# Structure and influence in an interconnected world: neurocomputational mechanism of real-time distributed learning on social networks

**DOI:** 10.1101/2022.03.22.485414

**Authors:** Yaomin Jiang, Qingtian Mi, Lusha Zhu

## Abstract

Many social species are embedded on social networks, including our own. The structure of social networks shapes our decisions by constraining what information we learn and from whom. But how does the brain incorporate social network structures into learning and decision-making processes, and how does learning in networked environments differ from learning from isolated partners? Combining a real-time distributed learning task with computational modeling, fMRI, and social network analysis, we investigated the process by which humans learn from observing others’ decisions on 7-node networks with varying topological structures. We show that learning on social networks can be realized by means similar to the well-established reinforcement learning algorithm, supported by an action prediction error encoded in the lateral prefrontal cortex. Importantly, learning is flexibly weighted toward well-connected neighbors, according to activity in the dorsal anterior cingulate cortex, but only insofar as neighbors’ actions vary in their informativeness. These data suggest a neurocomputational mechanism of network-dependent filtering on the sources of information, which may give rise to biased learning and the spread of misinformation in an interconnected society.

---

Social networks channel communication and route information transmission in human society^1,2^. By constraining what information we receive and from whom, the structure of social networks has substantial impacts on how we form beliefs and make decisions, and how collective opinion and behavior are shaped and propagated^3-5^. Although many studies have demonstrated the influence of social networks on political^6^, economic^7^, and social activities^8^ at the population level, the underlying neural and cognitive processes by which individuals incorporate information from connected peers remain to be explored. Answers to this question would shed light on the mechanisms of social learning and decision-making in a wider and more ecologically-relevant context, and help understand collective maladaptation––such as herding^9^ and misinformation propagation^10^––in terms of the computational challenges faced by individuals trying to process entangled information in an interconnected society.

A natural candidate for investigating learning on social networks from a neurocognitive perspective is the computational framework of reinforcement learning (RL)^11^. RL theories have been highly successful in connecting the cognitive and neurobiological bases with a broad range of behaviors^12-14^, including learning from social partners^15-20^. Despite its successes, the standard RL framework provides an incomplete account of learning in complex, interconnected environments. Consider, for example, the classic observational learning task^20-23^, where an individual learns from the decisions of multiple observees performing the same task as the observer. Prior RL-based research has typically assumed that the actions of different observees constitute independent learning signals, which can be statistically aggregated as an unbiased estimator for the common, unknown state of the environment for the observer^20,23,24^. Contrary to this assumption, however, substantial evidence in the social network literature suggests that choices selected by connected peers are often interrelated and vary in their informativeness^25,26^. Blindly relying on the conventional RL strategy without considering the underlying connections that spread social influences is essentially ignoring the potential variations and repetitions in social signals and can be detrimental to adaptive behavior in an interconnected environment^27^. Nevertheless, extant data suggest that social animals embedded in complex interaction webs demonstrate some level of sensitivity to the topological features of their immediate social environments^28,29^, leaving open whether and how relevant structural information is incorporated into the learning processes.

Theoretically, the social learning literature has proposed two classes of models to address the network effect. Normative strategies, such as Bayesian learning, assume that individuals rationally use the knowledge of the network structure to optimally distinguish between learning signals, filter out potential correlations in those signals, and integrate only the new piece of information into belief with the Bayes’ rule^7,30^. Despite its theoretical appeal, converging evidence suggests that Bayesian learning is cognitively unrealistic due to its excessive computational demand even on networks with relatively simple structures^30^. Naïve strategies, such as DeGroot learning, take an opposite, heuristic approach^31^. Although these models do not optimally adjust for signal heterogeneity and interdependency and sometimes lead to incorrect consensus among network members^7,30^, they provide simple quantitative accounts for how network geometry may affect learning. For instance, the now canonical DeGroot learning theory posits that learning on a network can be approximated by a Markov process driven by a weighted average of signals received from network neighbors. The weight used for signal aggregation reflects how strongly a particular individual is influenced by a neighbor, and has been linked theoretically to the network structure based on the limiting property of Markov processes^32^. However, no direct evidence is available for the bounded rational assumption at the heart of the naïve learning theories or for whether and how the underlying neurocognitive operations related to social influence are affected by network structures in ways that translate to trial-by-trial (rather than asymptotic) behavior^30,33^.

We hypothesized that the process by which the brain learns from networked environments could be characterized by incorporating the DeGroot heuristics into the RL framework. In the context of learning observationally from others’ decisions, the proposed DeGroot-RL model stands on the following three hypotheses that together enable us to delineate the rich network interactions in a quantitative yet neurobiologically-plausible manner. First, similar to the previous temporal difference algorithms of RL widely used in nonnetworked settings^11^, the DeGroot-RL model assumes that the brain integrates information across social observations by maintaining and updating an internal expectation signal. Learning from an action of a particular neighbor is driven by an action prediction error (aPE) between the observed and expected choice of this neighbor, weighted by a learning rate.

Second, the networked environment affects learning by differentially modulating the learning rate. Motivated by prior data on social influence that a well-connected individual has greater influence on her peers and is less susceptible to others’ opinion^34^, the DeGroot-RL model posits that the extent to which one learns from an observed action scales with the observee’s network connectedness, relative to that of the observer. Under this assumption, the brain needs to flexibly adjust the learning rate based on network locations, possibly according to signals related to the observee’s and observer’s degree centralities (i.e., the number of individuals to whom one is directly connected on a network), one of the most fundamental metrics for local prominence and immediate influence in social network analysis^2^.

The third hypothesis, derived from the DeGroot heuristics, postulates that the degree-modulation effect on learning may vary systematically over the course of information circulation, contingent on whether social observations differ in their informativeness^30^. For example, when individuals learn from others' firsthand, isolated information, the DeGroot-RL model will reduce to the standard RL-like algorithm for observational learning^21,22^, whereby an observer is equally influenced by the received information regardless of the differences in observees’ network locations or properties. In contrast, when learning from others’ secondhand, possibly heterogeneous and intertwining information, the strength of learning will be modulated by the relative degree centrality between the observee and the observer on the network. Under this hypothesis, the brain needs to process network-related information flexibly, according to its relevance to learning.

To assess these hypotheses and elucidate the neural and cognitive process related to the proposed DeGroot-RL algorithm, we used functional magnetic resonance imaging (fMRI) in conjunction with a distributed learning task for observational learning^30,35,36^ that was adapted from economic studies of information cascade^37^ and housed in a variety of exogenously given, 7-node, undirected, and unweighted networks. This task is simple enough to carry out in a controlled fMRI experiment with a reasonably large number of individuals interacting with one another in real time, yet it retains important features of opinion adaptation under social influence. Moreover, rather than focusing on a handful of special networks as in prior behavioral experiments^3,30^, we investigated the proposed model on a relatively large variety of network structures that were preselected based on the separability of choice behavior simulated by different learning models (*Methods*).

## Results

### A distributed learning game

A total of 217 unique subjects (31 fMRI participants) participated in the experiment in groups of 7 (1 inside the fMRI scanner; 209 included in data analyses with 25 fMRI participants; see *Methods* for subject exclusion). The experiment consisted of 40 separate games on varying networks (Extended Data Figs. 2-3; *Methods*). In each game, a participant’s goal was to infer an unknown state of the environment, which was common to all 7 participants in the game. At the beginning of a game, 7 subjects were randomly assigned to different nodes on a new network, and a computer selected one of two underlying states at random. Each subject received a private signal that was independently and identically distributed conditional on the same selected state, and needed to make an initial guess about the underlying state (Fig. 1A and Extended Data Fig. 1; *Methods*). We hypothesized that a subject should rationally base her estimation on the private signal in this decision. The prediction was confirmed by our data, in which 98.34 ± 5.12% (mean ± intersubject SD) initial estimations matched subjects’ private signals.

**Fig. 1.**
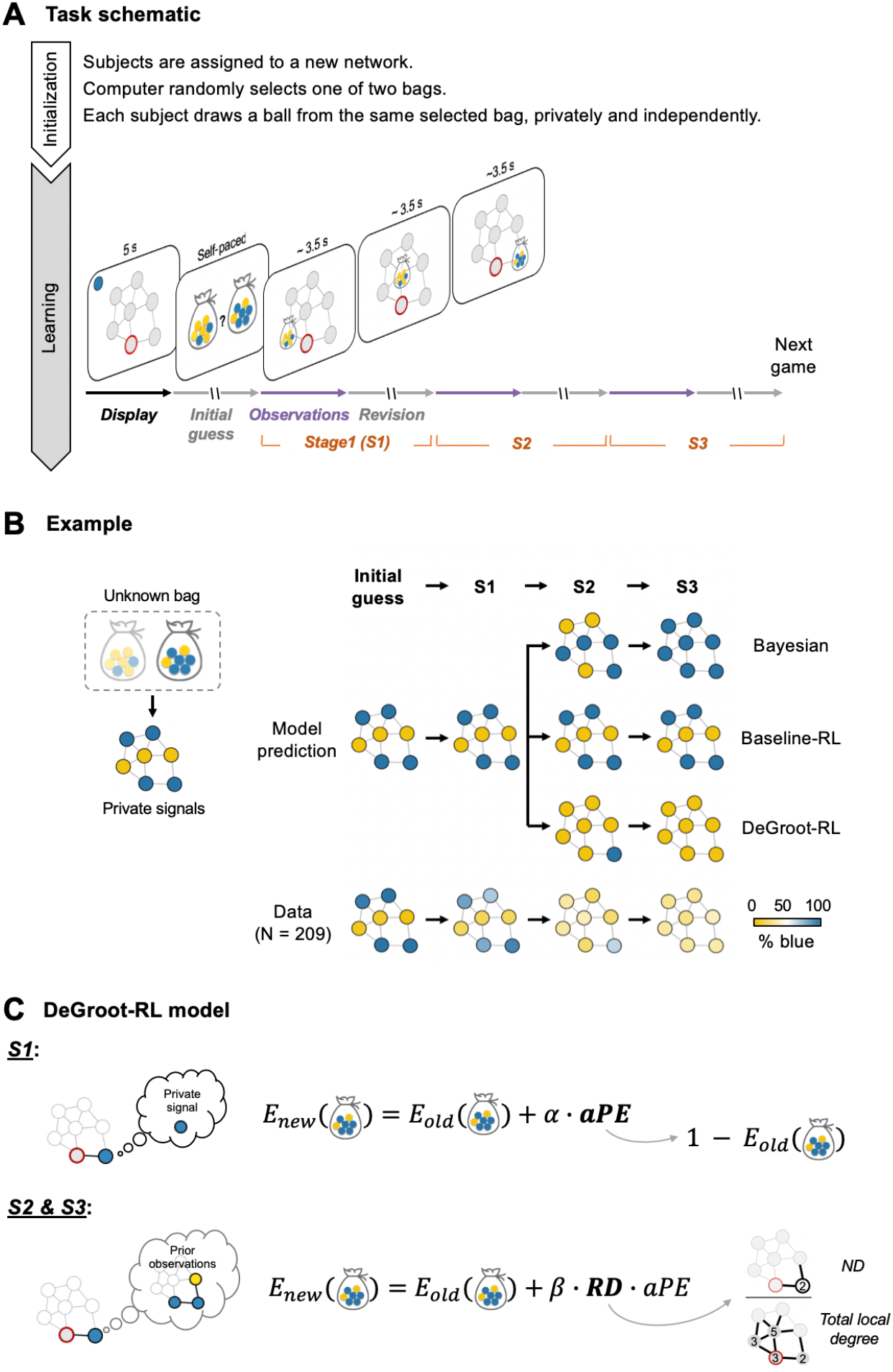
Distributed learning game and the DeGroot-RL model. **(A)** Task schematic. Each distributed learning game is initialized by randomly assigning subjects to different nodes on a new network and selecting one of two bags that contain yellow and blue balls with opposing ratios (5:2 vs. 2:5). Each subject, who does not know which bag was selected, privately draws a ball from the same selected bag with replacement, and needs to infer whether the underlying bag contains more blue or yellow balls in a series of decisions. All subjects are instructed that the chance of drawing a blue/yellow ball is either 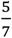 or 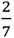, independently and identically distributed across all members embedded on the same network. After initialization, subjects are presented with the structure of the network (common for all network members; *Display*), one’s own network location (red circle in *Display*), and the ball privately drawn from the bag (top left corner in *Display*). Subjects are then simultaneously asked to decide between two candidate bags (*Initial guess*), followed by three stages of observational learning (*S1–S3*). At each stage, a participant is presented with the most recent decisions by her immediate neighbors (*Observations*) and then provided with an opportunity to reassess her previous decision (*Revision*). To facilitate the visual tracking of neighbors’ choices on a network display while counterbalancing the influence of observation order, neighbors’ choices are revealed sequentially, in a clockwise order, starting from a randomly-selected neighbor that varies across stages and between subjects. All 7 participants played the same game in real time from their respective network locations, with no feedback on the choice accuracy during the experiment (*Methods* and Supplementary Video 1). **(B)** Learning dynamics on an example network, which also serves to illustrate possible misinformation propagation on the network. Left: the underlying bag selected by the computer and the private signals given the selected bag. Right: Simulated and actual choices on the network. Color in each node represents a simulated choice (model prediction) or the frequency of actual choices (data) over bags containing more blue or yellow balls. While the Bayesian learning, baseline-RL, and DeGroot-RL models make similar predictions in the initial guess and S1 estimation, they differ sharply in S2 and S3 predictions, with the DeGroot-RL predictions best aligned with the actual choice frequencies of all participants on this network. Notably, this example also demonstrates a scenario in which network members reach an incorrect consensus under the DeGroot-RL strategy. Unlike rational Bayesian learners, who gradually form a consensus on the correct underlying state, both the simulated DeGroot-RL agents and actual participants converged toward the wrong estimation, biased by the inaccurate information from the central, most-connected individual on the network (see also Extended Data Fig. 4 for model simulation on all networks in the study). **(C)** DeGroot-RL model illustration. Left: An example illustrating the stage-varying information in an observed action. While an S1 observation reflects the neighbor’s private signal, an observation in S2 or S3 additionally signals what the neighbor has previously learnt from her neighbors. Right: Stage-dependent, degree-modulated learning. Upon observing an action from a neighbor, the belief expectation about the unknown state (*E*_*old*_) is updated through an action prediction error (*aPE*), defined as the discrepancy between the observed and expected action. Learning in S1 follows the typical RL setup where the aPE signal is scaled by a baseline learning rate (*α*). In S2 and S3, however, aPE signals are weighted by the learning rate (*β*) and relative degree (RD), with the latter being defined as the degree centrality of the observee relative to the total degree of the observer and all her direct neighbors on the network (*Methods*). Variants of DeGroot-RL formulations such as alternative learning rate specifications and RD definitions were evaluated against the proposed model in *Methods*.

Critical for the purpose of this study, the participants were then allowed to revise their estimations in response to the choices previously selected by the neighbors to whom each was directly connected on the network. To allow for meaningful fMRI analyses, a subject was presented with her direct neighbors’ prior decisions sequentially, one at a time, such that her neural responses could be directly linked to the action and network location of a particular observee (Fig. 1A and Extended Data Fig. 1; *Methods*). To allow for examining learning effects, the process of observing neighbors’ actions and reassessing one’s estimation was carried out 3 times consecutively within each game (henceforth *Stages 1-3*, or *S1*, *S2*, and *S3*; Fig. 1A).

Of note, 7 participants played the same game simultaneously, from their respective network locations via an intranet. That is, the participants were facing the same underlying state, the same network structure and display, and were making decisions at the same time in each game. Crucially, when a participant was witnessing her neighbors’ choices, her neighbors were also presented with the choice information from their respective neighborhoods (see illustration in Supplementary Video 1). Under such a real-time distributed learning setup, information received by an observer is incorporated into the observer’s subsequent decision and propagated gradually from the observer to her direct and indirect contacts along network connections in the later stages of the game. Throughout the experiment, all subjects were financially incentivized to guess as accurately as possible in all 4 decisions in each game (i.e., initial guess and 3 reassessments) and had no incentive to mislead or collude with others. No feedback was provided on the accuracy of estimations during the experiment, so that the only information a subject could rely on in a game was her private signal and the actions by her direct neighbors on the network (*Methods*).

This real-time, distributed learning game has two properties important for evaluating the DeGroot-RL model at the neurocomputational level. First, the three learning stages (S1–S3) were set up identically in each game (but with randomized observation order to control for potential order effects; *Methods*). This feature allowed us to evaluate the first DeGroot-RL hypothesis that subjects learned from neighbors’ actions in an error-driven manner and did so consistently across S1, S2, and S3. Second, despite their identical experimental setups, three learning stages differed in the type of information contained in neighbors’ actions. Unlike S1 observations, which reflect observees’ independent private signals, an S2 or S3 observation would additionally signal what the observee has learned from her neighbors (Fig. 1C, left), thereby becoming relatively more informative to the specific observer when the observee is better-connected and the observer is less-connected. Given this feature, the last two DeGroot-RL hypotheses led to a precise and testable prediction for the experiment––that is, learning from network neighbors might be modulated by the degree centrality of the observee relative to the observer in S2 and S3, but not in S1.

According to these predictions, we should expect not only the stage-dependent degree-modulation effect at the behavioral level, but also the differential involvement of neural signals of aPE and relative degree centrality––two core computational components in the DeGroot-RL model (Fig. 1C, right)––in the learning process. We predicted that there should be brain regions that track the aPE-related signals nonselectively in S1, S2, and S3, as well as brain regions that respond to signals associated with the relative degree centrality in S2 and S3, but not in S1.

### Learning behavior was modulated by degree centralities in S2 and S3, but not in S1

Behaviorally, participants in the experiment adapted their decisions in response to neighbors’ choices, such that the level of consensus within a network grew from 61.21 ± 1.29% (mean ± intergroup SD) in the initial guess to 88.10 ± 3.03% in the last (S3) decision, with no significant difference in the level of consensus changes across games between the early and late parts of the experiment (Pearson’s correlation between consensus change and game order: *r* = −0.05, *P* = 0.093; see also Extended Data Fig. 6 for behavioral dynamics in choice accuracy and more). To characterize the overall learning effect and evaluate DeGroot-RL predictions, we first performed model-free logistic regression analyses in each separate stage, to examine the extent to which the likelihood of participants altering their choices was influenced by social observations, and whether the strength of influence was modulated by the degree centralities of the observees and observers. We hypothesized that the likelihood that a participant aligned her estimation to an observation would be positively associated with the neighbor’s degree centrality (ND) but negatively associated with the observer’s own degree (OD) in S2 and S3, but not in S1.

To assess the impact of ND, we summarized the sequence of observations revealed to a subject at each stage of each game, using two variables: (i) sum of observations across all direct neighbors (1, if an observed action differs from the observer’s previous choice; otherwise, −1), and (ii) weighted sum of observations across all direct neighbors, with the respective neighbor’s degree serving as the weight. The inclusion of the unweighted and ND-weighted regressors helped to isolate the degree-modulation effect of interest from a baseline tendency of following the majority, a phenomenon widely reported in studies of group decision-making^9,38^. To remove any shared variances between the two regressors, we orthogonalized the ND-weighted regressor against the unweighted regressor, such that choices that were equally explainable by the two variables were attributed solely to the unweighted regressor. The regression coefficient for the ND-weighted sum of observations, therefore, served as a more stringent test on whether ND modulated learning.

Consistent with our hypothesis, mixed-effects regression conducted separately for each learning stage showed that the likelihood of a participant modifying her decision was positively correlated with not only the unweighted sum of observations, but also the ND-weighted sum of observations, in both S2 and S3 (Fig. 2A; sum of obs. in S2: *β* ± SEM = 3.26 ± 0.14, *z* = 23.40, *P* < 10^−15^; sum of obs. in S3, *β* = 2.58 ± 0.15, *z* = 16.91, *P* < 10^−15^; ND-weighted sum in S2: *β* = 1.51 ± 0.10, *z* = 15.68, *P* < 10^−15^; ND-weighted sum in S3: *β* = 1.22 ± 0.09, *z* = 14.10, *P* < 10^−15^). The positive effects suggested that, in addition to following the majority, subjects were more likely to be swayed toward the decisions of highly connected neighbors, relative to those of poorly connected neighbors embedded on the same network. In stark contrast, in S1, the ND-weighted regressor showed no extra explanatory power above and beyond the unweighted regressor in predicting participants’ subsequent choices (Fig. 2A; sum of obs. in S1, *β* = 3.26 ± 0.19, *z* = 16.81, *P* < 10^−15^; ND-weighted sum in S1, *β* = 0.04 ± 0.06, *z* = 0.63, *P* =0.528; see also Supplementary Table 2A for full regression results).

**Fig. 2.**
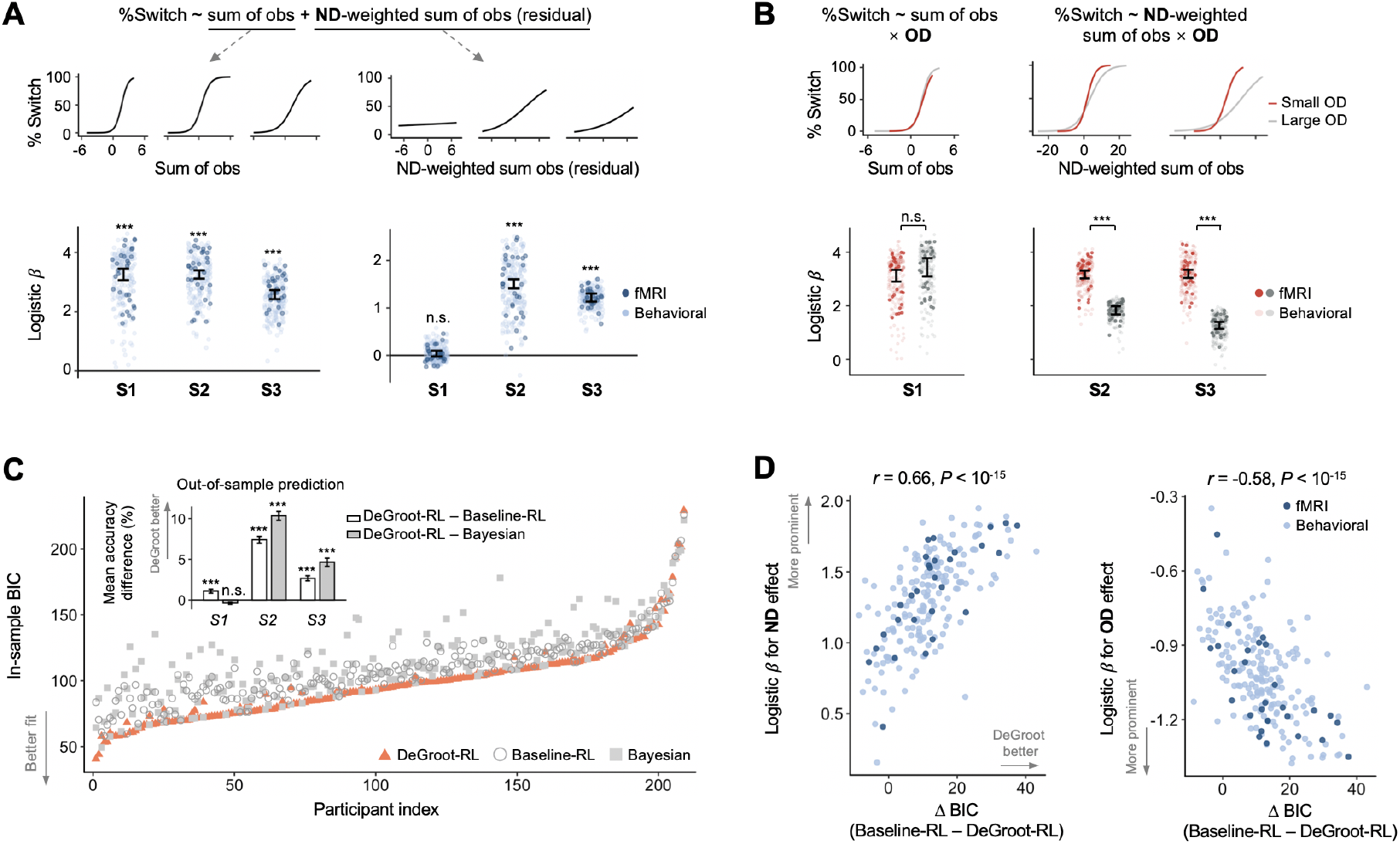
Behavioral evidence supporting the DeGroot-RL model. **(A)** Neighbor’s degree centrality (ND) modulates learning in S2 and S3, but not in S1. Logistic regression tests the probability of an observer modifying her previous estimation against the (unweighted) sum of observations (left) and the weighted sum of observations, with each respective neighbor’s degree serving as the weight (right). The ND-weighted regressor was orthogonalized against the unweighted regressor to remove shared variances. Top: the probability of changing a decision as predicted by the logistic regression estimates at each stage. Bottom: fixed-effects regression coefficients and SEM for the unweighted and ND-weighted sum of observations (depicted by error bars), overlaid by the random-effect coefficient associated with each observer (depicted by dots). Each dark/light dot represents an fMRI/behavioral participant. **(B)** Observer’s own degree centrality (OD) is negatively associated with the susceptibility to social observations in S2 and S3, but not in S1. The top and bottom panels show results from logistic regression testing the likelihood of changing one’s estimation in response to social observations, when an observer is highly vs. poorly connected. For the illustration purpose, large and small ODs were defined by median splits on OD across networks for each participant. **(C)** Comparisons of in-sample model fits using the Bayesian Information Criterion (BIC) of each participant show that DeGroot-RL explains behavioral choices better than Bayesian and baseline-RL (paired *t*-test, DeGroot-RL vs. baseline-RL, mean ± SEM = −11.58 ± 0.68, *t*_208_ = −17.07, *P* < 10^−15^; DeGroot-RL vs. Bayesian, −16.02 ± 1.14, *t*_208_ = −14.05, *P* < 10^−15^; see also Extended Data Fig. 5 for Bayesian model selection). Participants are sorted by the BIC score of the model that best explains choices. Inset: Out-of-sample prediction accuracy is superior for the DeGroot-RL model compared with Bayesian and baseline-RL models in S2 and S3. The observed difference in the prediction accuracy in S1 between DeGroot-RL and baseline-RL likely reflects the fact that the model parameters governing S1 predictions (e.g., *α* and inverse temperature) were set to maximize the likelihood of choices across all stages (rather than S1 only) and thus were affected by model configuration and estimation in other stages. **(D)** Across-subject correlation between model-free and model-based results. Individual BIC differences between DeGroot-RL and baseline-RL (*x*-axis) are plotted against the individual model-free estimates for ND (left) and OD (right) effects, respectively, exploiting the fact that the DeGroot-RL model differs from the baseline-RL model only in the assumption regarding the degree-modulation effect in S2 and S3 (see model setup in Fig. 1C and *Methods*). The model-free ND effect (*y*-axis, left) is captured by the coefficient of individual random effects with respect to the ND-weighted sum of observations (as shown by the dot in the bottom-right panel in Fig. 2A), averaged across S2 and S3 within each subject. The OD effect (*y*-axis, right) reflects the coefficient of individual random effects for OD × ND-weighted sum observations (as shown by the within-subject difference between the orange and grey dots in the bottom-right panel in Fig. 2B), averaged across S2 and S3 within each subject. A more negative OD effect (y-axis, right) corresponds to a more prominent reduction in the susceptibility to social influence as OD increases. These correlations remained significant when analyzing the across-subject association for S2 and S3 separately (Pearson’s correlation between ΔBIC and ND effect in S2, *r* = 0.62, *P* < 10^−15^, in S3, *r* = 0.32, *P* < 10^−5^; correlation between ΔBIC and OD effect in S2, *r* = −0.62, *P* < 10^−15^, in S3: *r* = −0.14, *P* = 0.039). Consistent results were also found after partialing out the variance shared between the model-free OD and ND estimates (partial correlation between ΔBIC and ND effect, *ρ* = 0.44, *P* < 10^−10^; ΔBIC and OD effect, *ρ* = −0.25, *P* < 0.001). Error bars represent SEM. *** *P* < 0.001, n.s., not significant, Bonferroni-corrected when appropriate.

To assess how learning was biased by a participant’s OD, we further compared the influence of the ND-weighted sum of observations on the observer’s subsequent decision when the observer was endowed with high vs. low degree centrality. Specifically, we carried out separate mixed-effects logistic regression for S2 and S3, testing the probability of an observer modifying her decision against the following variables: OD, ND-weighted sum of observations, and the interaction between these two variables. These analyses showed positive main effects for both the OD and ND-weighted regressors (Supplementary Table 2C). More importantly, and consistent with the DeGroot-RL prediction, a negative interaction was seen between these variables in both S2 and S3 (Fig. 2B; interaction in S2: *β* = −1.10 ± 0.11, *z* = −9.69, *P* < 10^−15^; S3: *β* = −1.00 ± 0.09, *z* = −11.43, *P* < 10^−15^), suggesting a decreased susceptibility to social observations when a participant was highly relative to poorly connected. This result was not due to the specific way of summarizing the sequence of observations within each stage and remained significant when examining the interaction between OD and the unweighted (rather than ND-weighted) sum of observations (Supplementary Table 2B). In S1, by comparison, no systematic variation was observed in participants’ susceptibility to neighbors’ actions with their OD, as revealed by a similar regression analysis for the likelihood of altering one’s decision against variables of OD, unweighted sum of observations, and the interaction between the two (Fig. 2B; interaction in S1: *β* = 0.15 ± 0.17, *z* =0.92, *P* = 0.357; see also Supplementary Table 2B).

### The DeGroot-RL model characterized learning behavior better than alternative models

To more formally test the DeGroot-RL predictions and derive latent variables that might reflect neurocognitive operations underlying learning on networks, we fit the DeGroot-RL model with each participant’s choice behavior (*Methods*). As shown in Fig. 1C, our proposed DeGroot-RL model assumed that the S2 and S3 learning rate was modulated by a measure of relative degree (RD), defined as the observee’s degree with respect to the total degree of the observer and all her direct neighbors on the network (*Methods*). The normalization term in RD reflected the mathematical requirement that a learning rate should be no greater than 1, although the alternative specification using the observer’s degree as the denominator yielded similar results (*Methods* and Supplementary Table 1).

We compared the proposed DeGroot-RL model against two benchmark models: a baseline-RL model, which assumed that the network-related information was completely ignored throughout learning, and a Bayesian learning model, which assumed that the information regarding the network structure was rationally used by all participants (*Methods*). It is worth noting that the DeGroot-RL model differed from the Bayesian and baseline-RL models in predicting choices mainly in S2 and S3, but was similar to those models in S1, such that all three models predicted the unweighted, rational integration of S1 observations^30^ (*Methods*; see also Fig. 1B for an example).

We computed both the Bayesian Information Criterion (BIC) based on the in-sample model fit and out-of-sample prediction accuracies using a five-fold cross-validation procedure (*Methods*; see also Supplementary Table 3 for model estimation results). Both measures showed that the DeGroot-RL model outperformed the alternative models. In particular, the DeGroot-RL model had a lower BIC (better fit) than the alternative models (Fig. 2C; DeGroot-RL = 20407.41; baseline-RL = 22827.97; Bayesian = 23755.10), with its BIC score being the lowest in more than 78% of participants (164 out of 209 participants; see also Extended Data Fig. 5 for Bayesian model selection). Based on the prediction accuracy computed from holdout samples, the DeGroot-RL model predicted subjects’ behavior with an accuracy of 75.29 ± 0.62% (mean ± intersubject SEM, averaged over all four decisions within each game), which was significantly higher than that from the baseline-RL model (paired *t*-test, *t*_208_ = 15.38, *P* < 10^−15^) and the Bayesian learning model (*t*_208_ = 15.87, *P* < 10^−15^; see also Extended Data Fig. 5). This increase in the prediction accuracy was largely contributed by the improved prediction in both S2 and S3, rather than an enhancement coming from either S2 or S3 alone (inset, Fig. 2C).

Importantly, choice simulation based on the DeGroot-RL model estimates could successfully recover the model-free patterns in each learning stage (Extended Data Fig. 7). In contrast, simulations from the Bayesian and baseline-RL models––even with the best-fitting parameters calibrated on choice behavior––failed to capture key behavioral features in S2 and S3, such as the ND and OD effects revealed by the logistic regression analyses of the actual data (Extended Data Fig. 7). Besides the aggregate choice patterns, the DeGroot-RL model estimates were also consistent with the model-free analyses at the across-subject level, such that subjects whose behavior was better characterized by DeGroot-RL showed a more pronounced behavioral sensitivity to both ND and OD in S2 and S3, as measured by the respective logistic regression estimates for individual participants (Fig. 2D).

Besides individual choices, the proposed DeGroot-RL model also captured the variations in other behavioral measures, including choice difficulty as reflected by participants’ reaction time, the dynamics of choice consensus, and the trajectory of estimation accuracy (Extended Data Fig. 6). In addition to the Bayesian and baseline-RL models, the proposed model also outperformed a range of alternative models that could account for participants’ choices based on assumptions differing in either which network measure (rather than RD) might modulate learning or how RL parameters were specified in the model (*Methods* and Supplementary Table 1). Finally, to address the potential concern regarding the between-network variations in learning behavior, we conducted model estimation for each network (pooling over subjects), and observed similar results using both the in-sample and out-of-sample measures for goodness-of-fit at the across-network level (*Methods* and Extended Data Fig. 5).

### Right lateral prefrontal cortex (rLPFC) tracked aPE estimates in S1, S2, and S3

Having established the DeGroot-RL model at the behavioral level, we then investigated whether fMRI activity reflected key computational components of the model, including aPE and RD, on an observation-by-observation basis, at the time when participants were presented with a neighbor’s action. We also tested whether the neural responses to the aPE and RD signals would demonstrate differential stage-related patterns, as predicted by the DeGroot-RL model. We conducted a standard general linear model (GLM) analysis on fMRI data, entering the value estimate of aPE associated with each observed action derived from the best-fitting DeGroot-RL model for each individual, together with the RD value associated with the respective observee and observer, at the observation onsets of the corresponding learning stage (GLM1, *Methods*). Parametric regressors were orthogonalized against one another in all GLM analyses in the present study, such that the regression coefficients captured the variations in blood-oxygen-level-dependent (BOLD) signals in the specific brain regions that were uniquely explained by each regressor, rather than the shared variances.

As we were particularly interested in evaluating whether there were any brain regions consistently tracking aPE estimates across three learning stages, we averaged the GLM1 coefficients of the aPE estimates across stages for each subject before taking them into the group-level analyses. This identified significant neural responses in the rLPFC (Fig. 3A; see also Supplementary Table 5 for full activation including the middle temporal gyrus and visual cortex), which has been previously implicated in representing notions of prediction error signals in action observation learning^21,22,24^ and as part of the “mirror” system encoding the executed and observed actions in a range of interpersonal scenarios^39^. Rather than being driven by a simple effect such as the correlation with the observed action, activity in the identified rLPFC cluster reflected both computational components in an aPE signal: It scaled positively with the observed action (1 if the observation matches the observer’s previous decision; *β* = 0.23 ± 0.04, *t*_24_ = 5.29, *P* < 0.0001; Extended Data Fig. 8), but negatively with the observer’s expectation about the underlying state (*β* = −0.12 ± 0.06, *t*_24_ = −2.23, *P* = 0.036; Extended Data Fig. 8). As a robustness check, we tested additional decision variables that might be related to the processing of social observations, including color selected by the observee (yellow/blue), order in which the neighbor’s action was presented in the particular stage of the game, and the consensus level among network members at the beginning of the stage (GLM2, *Methods*). The observed rLPFC encoding could not be attributed to any of these variables and remained significant with the inclusion of these variables as regressors of no interest in one regression model (cluster-wise family-wise-error(FWE)-corrected *P* < 0.05, with cluster-forming threshold uncorrected *P* (*P_unc._*) < 0.001).

**Fig. 3.**
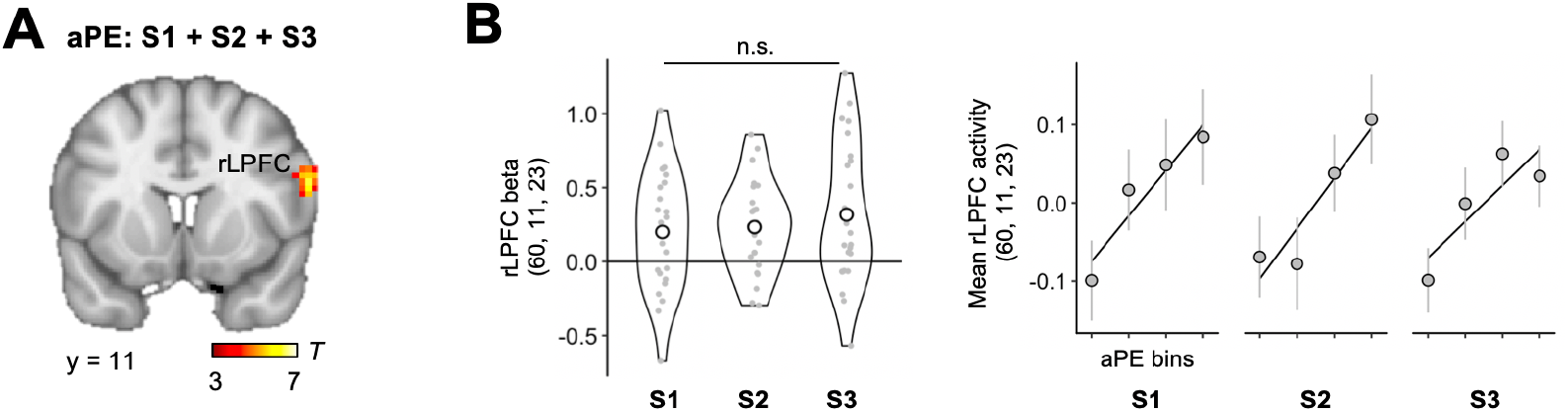
Right lateral prefrontal cortex (rLPFC) tracks the value estimate of action prediction error (aPE) in S1, S2, and S3. **(A)** Statistical parametric map with respect to aPE estimates at observation onsets, computed by averaging the GLM1 coefficients for aPE estimates across S1, S2, and S3 for each subject and then taking them into the standard random-effects group analysis (peak voxel Montreal Neurological Institute (MNI) coordinates: *x*, *y*, *z* = 60, 11, 23; cluster-wise family-wise-error(FWE)-corrected *P* < 0.05, with cluster-forming threshold *P*_*unc.*_ < 0.001). **(B)** rLPFC cluster demonstrates similar effect sizes for individual aPE estimates across stages. Left: Violin plots for the distribution of individual rLPFC beta values for aPE estimates, separately extracted for each stage from the significant rLPFC cluster as identified in Fig. 3A. Each dot represents a subject. Each circle represents a group mean. Right: Mean fMRI activity extracted from the same rLPFC cluster and binned by aPE estimates in each separate stage. Similar results were obtained by a whole-brain paired comparison, showing that no significant cluster differentially responded to aPE estimates across stages at cluster-wise FWE-corrected *P* < 0.05. N.s., not significant. Error bars represent intersubject SEM.

Moreover, the rLPFC cluster stably tracked aPE estimates across S1, S2, and S3, demonstrating similar effect sizes across stages in the regression estimates (beta values) that were extracted from the significant rLPFC cluster for each separate stage (Fig. 3B, left; one-way repeated-measures ANOVA, *F*(2,48) = 0.55, *P* = 0.582). To illustrate this result, we also plotted the mean activity obtained from the rLPFC cluster as a function of four bins of ascending aPE estimates in separate learning stages and observed similar correlation patterns (Fig. 3B, right). This result was further confirmed by a whole-brain within-subject comparison of the aPE correlates, in which we identified no significant cluster that responded differentially to aPE estimates across stages at cluster-wise FWE-corrected *P* < 0.05.

In addition to aPE signals at observation onsets, we also tested for brain regions responding to other classic learning signals at the time of decision submission in each learning stage. Consistent with prior neuroimaging evidence, we observed activity in the orbitofrontal cortex (OFC) signaling the reward expectation estimate associated with the chosen option^13,14^, as well as activity in the anterior cingulate cortex (ACC) and the neighboring medial prefrontal cortex reflecting the model-derived probability of switching away from one’s prior estimation^40^ (Extended Data Fig. 9). Findings at the choice time, together with the aPE signals at the observation time, indicated an error-driven process similar to the temporal difference form of RL during action observation learning channeled by social networks.

### Dorsal anterior cingulate cortex (dACC) represented RD-related signals in S2 and S3

Next, we tested whether activity in any brain regions would reflect the observation-by-observation changes in the relative degree centrality between the observee and the observer (i.e., RD) at observation onsets, and did so stably across S2 and S3. Similar to the analysis of aPE signals, we first looked for the RD correlates by averaging the GLM1 coefficients for RD over S2 and S3 for each fMRI participant (*Methods*). This revealed a strong correlation in a network of brain regions, including the dorsal anterior cingulate cortex (dACC) extending to the adjacent presupplementary motor area (preSMA), precuneus, bilateral anterior insula, visual cortex, and other areas (Fig. 4A; see also Supplementary Table 5 for full activation list). The loci of activation in the dACC/preSMA were similar to those seen in the past experiments where subjects adjusted behavioral strategies, such as learning rate, in response to environmental changes^41-45^. Moreover, dACC activity demonstrated features consistent with the assumption that the relative rather than absolute value of degree centrality was involved in learning. Activity in the dACC simultaneously correlated with the neighbor’s degree (numerator in RD) and total local degree (denominator in RD), with opposing signs, at observation onsets in both S2 and S3. Specifically, this opposing correlation pattern was observed not only within a region of interest (ROI) in the dACC independently defined using an automated online meta-analysis^46^ (Fig. 4B, right), but also in a whole-brain conjunction analysis across positive activation for neighbor’s degree and negative activation for the total local degree in S2 and S3 (Fig. 4A).

**Fig. 4.**
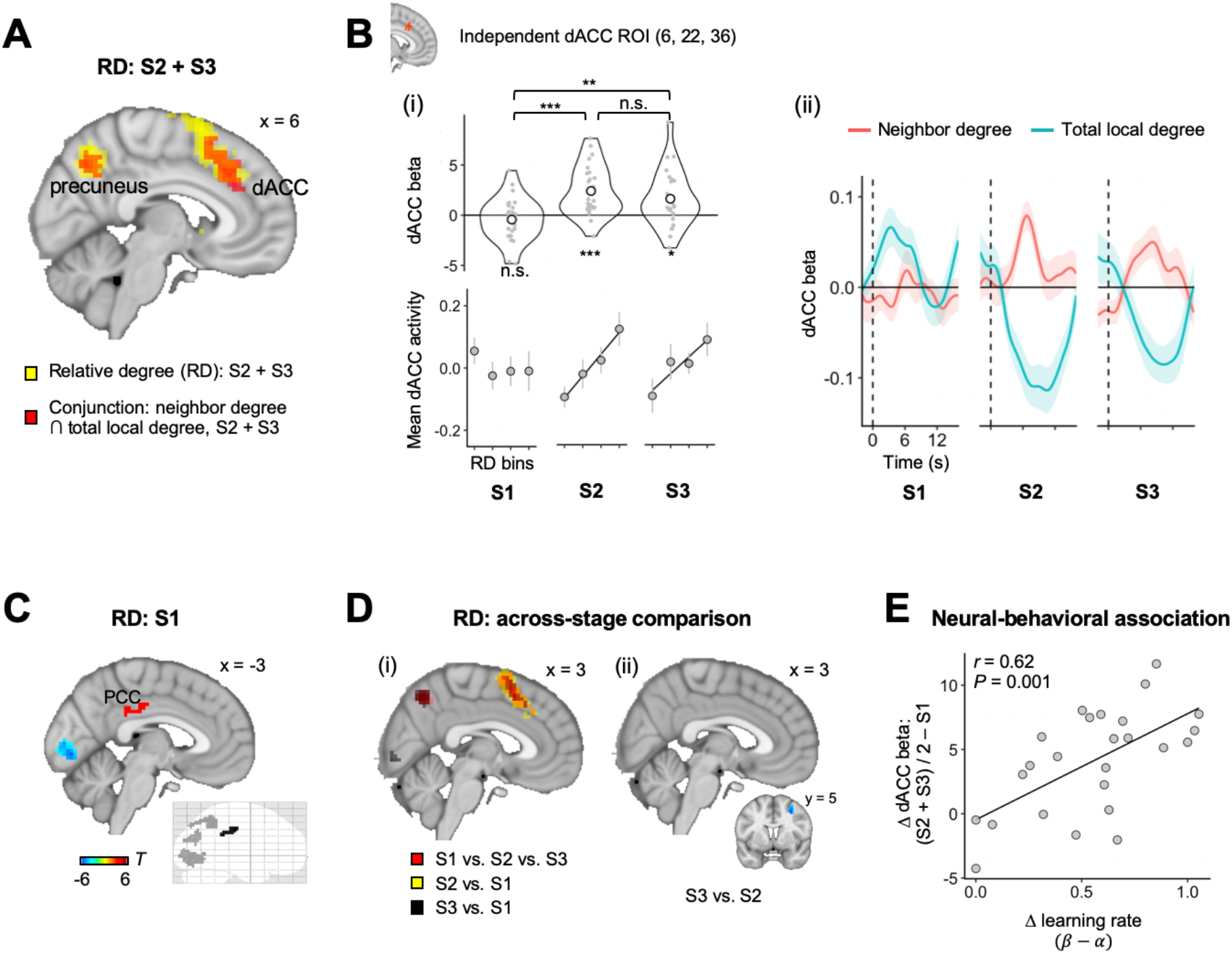
Activity in the dorsal anterior cingulate cortex (dACC) correlates with the value of relative degree (RD) in S2 and S3, but not in S1. **(A)** dACC shows RD-related signals on an observation-by-observation basis in S2 and S3. Regions shaded in yellow indicate clusters where activity significantly correlates with RD values at observation onsets in S2 and S3, calculated by averaging the GLM1 coefficients for RD across S2 and S3 for each participant and then taking them into the standard group-level analysis (cluster-wise FWE-corrected *P* < 0.05, with cluster-forming threshold *P*_*unc.*_ < 0.001; see also Extended Data Fig. 10A for the extent of RD encoding). Regions shaded in red indicate clusters scaling both positively with neighbor’s degree (numerator in RD) and negatively with total local degree (denominator in RD) at observation onsets in S2 and S3, as revealed by a whole-brain conjunction analysis for overlapping activation between the neighbor’s degree and total local degree (cluster-wise FWE-corrected *P* < 0.05, with cluster-forming threshold *P*_*unc.*_ < 0.001; *Methods*). The conjunction result was computed by first averaging the individual GLM coefficients for neighbor’s degree (or total local degree; GLM4, *Methods*) across S2 and S3 in the same way we looked for the RD correlates, and then using the resulting statistical maps to test for overlapping activation between either (i) the positive encoding of neighbors’ degree and negative encoding of total local degree (shown in Fig. 4A), or (ii) the negative encoding of neighbors’ degree and positive encoding of total local degree (no significant overlap at cluster-wise FWE-corrected *P* < 0.05; *Methods*). **(B)** dACC region of interest (ROI), independently defined by Neurosynth^46^. Top-left: beta values with respect to RD extracted from the same independent ROI at observation onsets in separate learning stages (S1: *β* = −0.43 ± 0.44, *t*_24_ = −0.99, *P* = 0.332; S2: *β* = 2.41 ± 0.50, *t*_24_ = 4.85, *P* < 10^−4^; S3: *β* = 1.64 ± 0.56, *t*_24_ = 2.91, *P* = 0.008; one-way repeated-measures ANOVA, *F*(2,48) = 10.65, *P* < 0.001). Bottom-left: mean dACC activity binned by RD values in each stage. Error bars represent intersubject SEM. Right: Time-course analyses with respect to the neighbor’s degree and total local degree for each stage within the same independent dACC ROI. Vertical dashed lines indicate the observation onset. **(C)** Neural correlates of RD values at observation onsets in S1 (cluster-wise FWE-corrected *P* < 0.05, with cluster-forming threshold *P*_*unc.*_ < 0.001; see also Extended Data Fig. 12 for ROI analyses in S1). **(D)** Paired comparisons with respect to RD correlates across learning stages. Left: Clusters shaded in red show results of whole-brain ANOVA analysis comparing RD correlates across S1, S2, and S3 within subjects. Clusters in yellow and black show post hoc ANOVA analyses testing the stage effect by comparing the RD correlates in S2 vs. S1 (yellow) and S3 vs. S1 (black) (all cluster-wise FWE-corrected *P* < 0.05, with cluster-forming threshold *P*_*unc.*_ < 0.001). Right: Post hoc paired comparison of RD correlates between S2 and S3. The only significant cluster locates in the middle frontal gyrus (MNI: *x, y, z* = 30, 5, 47; cluster-wise FWE-corrected *P* < 0.05, with cluster-forming threshold *P*_*unc.*_ < 0.001). **(E)** Across fMRI subjects, the dACC beta values with respect to RD in S2 and S3, relative to that in S1, is positively correlated with the individually-estimated learning rate in S2 and S3, relative to that in S1. This effect was significant not only when we averaged the individual’s dACC beta over S2 and S3 (Pearson’s *r* = 0.62, *P* = 0.001), but also when we tested the effect in S2 and S3 separately, with no significant difference between these stages (S2: *r* = 0.52, *P* = 0.007; S3: *r* = 0.53, *P* = 0.006; S2 vs. S3: *β* = −0.11 ± 0.25, *t*_24_ = 0.43, *P* = 0.673). Each dot represents a subject. ^*^ *P* < 0.05, ^**^ *P* < 0.01, ^***^ *P* < 0.001, n.s.: not significant, Bonferroni-corrected when appropriate.

Analyses further showed that the RD values explained dACC activity at observation onsets above and beyond a range of decision-related variables, including visual properties associated with the network display, nuisance effects arising from action observation, and cognitive components that might be related to learning or to other dACC functions implicated by previous studies. In particular, we performed another GLM analysis (GLM3, *Methods*), which additionally included the observee’s visual centrality in the network display (i.e., the Euclidean distance between the observee’s location and the visual center of the layout), the visual distance between the observee’s and observer’s location in the network display, order of observation display, color selected by the observee (yellow/blue), magnitude of aPE estimates, variance in attained observations within the current stage, level of conflict between social observations and the observer’s decision (i.e., the proportion of attained observations within the current stage that were different from the observer’s prior decision), updated belief expectation associated with the observer’s prior decision, and choice difficulty reflected by the distance in the belief expectation estimates between two choice options. None of these variables could explain the same portion of dACC activation as RD values and the observed parametric encoding of RD remained significant in the dACC, even after regressing out the influence of all these variables as regressors of no interest in the same GLM model (cluster-wise FWE-corrected *P* < 0.05, with cluster-forming threshold *P*_*unc.*_ < 0.001; Extended Data Figs. 10-11).

Moreover, activity in the dACC demonstrated similar response patterns to the RD values in S2 and S3. Neural betas separately extracted for S2 and S3 from the same independent dACC ROI were both highly significant and showed no systematic difference in their effect sizes (Fig. 4B; S2: *β* = 2.41 ± 0.50, *t*_24_ = 4.85, *P* < 10^−4^; S3: *β* = 1.64 ± 0.56, *t*_24_ = 2.91, *P* = 0.008; S2 vs. S3: *β* = 0.77 ± 0.72, *t*_24_ = 1.07, *P* = 0.294). This result was further confirmed by a whole-brain within-subject comparison of RD correlates. Except for a cluster confined to the middle frontal gyrus (MNI: *x, y, z* = 30, 5, 47; Fig. 4D, right), we found no other cluster that responded differently to RD values in S2 vs. S3 at cluster-wise FWE-corrected *P* < 0.05 (Fig. 4D, right).

### Activity in the dACC did not correlate with RD values in S1

By contrast, in S1, the same GLM1 analysis revealed no significant correlation with RD values at observation onsets in the dACC or other frontal regions, in either positive or negative direction (Fig. 4C; cluster-wise FWE-corrected *P* < 0.05, with cluster-forming threshold *P*_*unc.*_ < 0.001). Instead, we observed positive correlations with RD values in a circumscribed cluster in the posterior cingulate cortex (PCC), and negative correlations restricted to the precuneus and visual cortex (Fig. 4C and Extended Data Fig. 12). The identified regions in the PCC and precuneus have been recently implicated in encoding features of real-world social networks, even when such network features were task-irrelevant^29,47,48^.

To more formally examine the spatial expression of the RD correlates and test the stage-varying involvement of the dACC, we searched the whole brain for voxels that responded similarly (conjunction analyses) or differently (ANOVA analyses) to RD values across stages. A three-way conjunction among S1, S2, and S3 showed no significant cluster for either positive or negative activation to RD values (cluster-wise FWE-corrected *P* < 0.05, with cluster-forming threshold *P*_*unc.*_ < 0.001; Extended Data Fig. 10). Importantly, this lack of overlapping activation could not be attributed to the lack of overlaps between S2 and S3, as an additional conjunction analysis between S2 and S3 identified substantial activation to RD, including a large cluster in the dACC, that considerably overlapped with the activation areas identified in Fig. 4A (Extended Data Fig. 10). Moreover, using a whole-brain ANOVA analysis, we directly compared RD correlates across stages within subjects, and identified a significant stage effect in several brain regions including the dACC (Fig. 4D, left; cluster-wise FWE-corrected *P* < 0.05, with cluster-forming threshold *P*_*unc.*_ < 0.001). As shown by post hoc paired comparisons, this stage difference was attributable to the increased correlation between dACC activation and RD values in S2 vs. S1, and S3 vs. S1 (Fig. 4D, left), but not by S2 vs. S3, in either positive or negative direction (Fig. 4D, right; see also Fig. 4B for ROI analyses). Together, these data provided consistent evidence suggesting that the neural correlates of RD in S1 were spatially segregated from those in S2 and S3 in a manner consistent with the DeGroot-RL prediction.

### dACC sensitivity to RD values was predictive of behavioral sensitivity to RD

To relate the encoding of RD values in the dACC to choice behavior, we tested whether, across subjects, the extent to which dACC activity reflected RD values was predictive of the behavioral effects of RD on learning. We used the individual value estimate of learning rate in S2 and S3 (i.e., *β* as in Fig. 1C) as a measure of how strongly RD affected learning at these stages (zero effects on learning when *β* = 0). To capture the overall individual neural sensitivity to RD, we averaged the dACC beta of RD from S2 and S3 in each subject. We then plotted the individual behavioral estimate of *β* against the dACC beta in S2 and S3, controlling for the respective baseline effects in S1 (Fig. 4E). The data showed that subjects with higher learning rates in S2 and S3 than in S1 exhibited greater dACC sensitivity to RD values at observation onsets in S2 and S3 than in S1 (Pearson’s *r* = 0.62, *P* = 0.001). This between-subject association not only held for dACC beta values averaged over S2 and S3, but was also highly significant and denoted similar effect sizes when tested separately in these stages (S2: *r* = 0.52, *P* = 0.007; S3: *r* = 0.53, *P* = 0.006; S2 vs. S3: *β* = −0.11 ± 0.25, *t*_24_ = 0.43, *P* = 0.673). Notably, this neural-behavioral association was not a spurious effect arising from the double dipping of data^49^, because the dACC beta of RD was purely determined by the neural responses to the exogenously-given networks, independent of participants choices, model specification, or data estimation.

### Ventromedial prefrontal cortex (VMPFC) signaled the value estimate of updated belief expectation at the time of observation in S1, S2, and S3

The above results thus raised the question of how social observations from disparate neighbors were integrated in the brain to inform the subsequent decision. Unlike previous learning experiments, where subjects typically make a choice immediately after an observation, our experiment required participants to cache a sequence of social information until they were asked to make a decision. Thus, a sensible strategy based on the DeGroot-RL hypotheses would be to maintain an expectation about the unknown state and sequentially update the expectation using either the unweighted (in S1) or RD-weighted (in S2 and S3) prediction error signals each time an observation is witnessed.

This hypothesis immediately led to two neural predictions. First, signals reflecting the value estimate for updated expectation (*E*_*new*_ as in Fig. 1C) might be represented in brain regions previously implicated in tracking RL expectations, like the OFC or VMPFC^13,14^. That is, in addition to the classic RL signals for belief expectations of the chosen option at choice time (as shown in Extended Data Fig. 9A), we would also expect––at observation times––the neural representation of *E*_*new*_ estimate associated with the option previously selected by the observer that has been updated according to the observed action. Similar to aPE signals, we hypothesized that signals related to *E*_*new*_ estimates would be seen on an observation-by-observation basis and across three stages nonselectively. The second prediction was motivated by the DeGroot-RL hypothesis that, compared to S1, incorporating an aPE signal into belief expectation in S2 and S3 would involve additional modulatory inputs. Thus, regions representing *E*_*new*_ estimates might demonstrate increased functional connectivity in S2 and S3 compared to S1, with regions related to tracking, representing, or implementing modulatory signals in service of learning.

We tested the first prediction in a new GLM (GLM5, *Methods*), which included the value estimate of *E*_*new*_ as a parametric modulator at the time of observation onset of in the corresponding learning stage. To control for decision factors that might be related to learning or belief updating, we included the following variables as regressors of no interest in the same regression model: aPE estimate associated with the observed action, RD, and the order of observation display. Similar to the above analyses for aPE estimates, we averaged each individual’s regression coefficients with respect to *E*_*new*_ estimates across three stages and then took them into second-level analyses. We found a strong positive correlation between *E*_*new*_ estimates and activity in a number of brain regions including the VMPFC (Fig. 5A; cluster-wise FWE corrected *P* < 0.05, with cluster-forming threshold *Punc*. < 0.001; Supplementary Table 5). The observed VMPFC responses to *E*_*new*_ estimates were robust (cluster-wise FWE-corrected *P* < 0.05, with cluster-forming threshold *P*_*unc.*_ < 0.001) to the inclusion of additional decision-related variables, such as the product of the RD value and aPE estimate, color selected by the observee, visual properties related to the network display (visual centrality and distance), and the global consensus level at the beginning of the learning stage (GLM6, *Methods*). As predicted, we observed stable neural representation of *E*_*new*_ estimates across S1, S2, and S3, such that within-subject comparisons identified no significant difference in neural responses to *E*_*new*_ estimates across stages either within the VMPFC cluster (Fig. 5B; one-way repeated-measures ANOVA, *F*(2,48) = 2.33, *P* = 0.108) or at the whole-brain level (cluster-wise FWE-corrected *P* < 0.05, with cluster-forming threshold *Punc*. < 0.001).

**Fig. 5.**
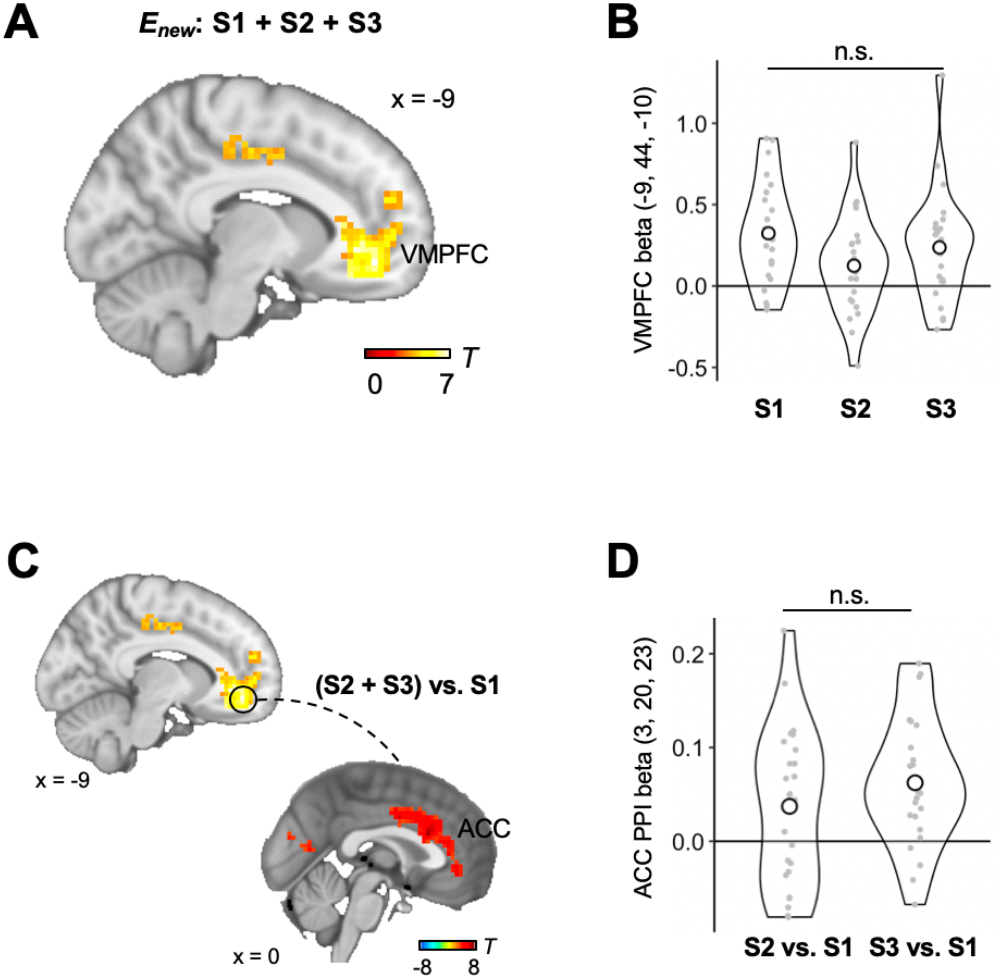
Ventromedial prefrontal cortex (VMPFC) tracks the value estimate of updated belief expectation (*E*_*new*_) in S1, S2, and S3. **(A)** Statistical parametric map with respect to *E*_*new*_ estimates at observation onsets, computed by averaging the GLM5 coefficients for *E*_*new*_ across S1, S2, and S3 for each subject in the same way we looked for neural correlates of aPE estimates (MNI: *x*, *y*, *z*, = −9, 44, −10; cluster-wise FWE-corrected *P* < 0.05, with cluster-forming threshold *P*_*unc.*_ < 0.001; *Methods*). **(B)** Similar effect sizes in the parametric encoding of *E*_*new*_ estimates in the VMPFC across stages, as demonstrated by violin plots for the distribution of neural betas for *E*_*new*_ estimates. The neural betas were extracted for each separate stage from the significant VMPFC cluster as identified in Fig. 5A. **(C)** Increased functional connectivity between the seed region in the VMPFC and a cluster in the anterior cingulate cortex (ACC) at observation onsets in S2 and S3, relative to S1 (cluster-wise FWE-corrected *P* < 0.05, with cluster-forming threshold *P*_*unc.*_ < 0.001). Similar to other GLM analyses in the study, this psychophysiological interaction (PPI) analysis compared the connectivity strength averaged over S2 and S3 against that in S1 (*Methods*). The seed region was defined as a 6-mm sphere around the peak activation as identified in Fig. 5A. **(D)** No systematic difference in the effect sizes of functional coupling between S2 and S3 (paired *t*-test, *t*_24_ = −1.20, *P* = 0.242), as revealed by the PPI betas extracted from the significant cluster in the ACC as identified in Fig. 5C.

To test the second prediction, we performed an exploratory psychophysiological interaction (PPI) analysis to look for brain regions that showed differential functional coupling with the VMPFC when RD was vs. was not needed for scaling aPE signals. Seeded in the VMPFC (6 mm around the peak activation as identified in Fig. 5A), the PPI analysis compared the connectivity strength averaged over S2 and S3 against that in S1 (*Methods*). This showed increased coupling of the VMPFC with several regions including a cluster in the ACC (Fig. 5C; cluster-wise FWE-corrected *P* < 0.05, with cluster-forming threshold *P*_*unc.*_ < 0.001), with no significant difference in the coupling effect sizes between S2 and S3 (Fig. 5D; paired difference for S2 – S3 = −0.03 ± 0.02, *t*_24_= −1.20, *P* = 0.242). The cluster identified by the PPI analysis partially overlapped with the area in the dACC signaling RD values in S2 and S3 (Extended Data Fig. 13), yet its peak activation was located in the more ventral and rostral portion of the ACC (MNI: *x*, *y*, *z* = 3, 20, 23; Fig. 5C). On the one hand, the overlapping dACC activation points to a possibility that the dACC might be involved in both representing the degree-related information and using this information for modulating the VMPFC representation, when network features are relevant for learning. On the other hand, as the ACC is a richly intra-connected system with projections to a broad set of regions, including the VMPFC^50^, it is also possible that the dACC cluster identified for encoding RD played an indirect role in influencing the VMPFC, through its effects on other parts of the ACC, such as its ventral and rostral portions, that have been previously implicated in monitoring and integrating learning signals^19,42^.

## Discussion

Information flowing in a large-scale, interconnected society is often entangled, conflated, and sometimes superfluous^1,26^. This poses a computational challenge for social learning, during which agents need to reconcile disparate sources of signals based on their informativeness^51^. Prior research on individual learning in non-social contexts has shown that humans can accurately estimate how relevant a learning signal is in predicting future and use this estimate to adjust RL learning rates^42,52,53^. On social networks, however, optimally evaluating the predictive value of each observation is cognitively demanding, sometimes even prohibitive. Indeed, failure to effectively aggregate information from connected peers has long been hypothesized to underlie herding, social influence biases, misinformation propagation, and other forms of collective maladaptation^9,54^.

Combining fMRI, formal theories of RL, and social network analysis, we explored the possibility that to balance computational costs, the brain approximates the relative informativeness of a social signal based on the structural properties of the network that routes information transmission. To evaluate this possibility, we grounded the DeGroot learning heuristics, a classic theory for naïve social learning imported from social network analysis^7,30,31^, into the temporal difference form of RL, widely implicated in the neurobiology of learning and decision-making^21,42,52^. Using a real-time, distributed learning task on networks with varying topological structures, this study provided behavioral and neural evidence that learning in complex, interconnected environments can be realized by means similar to the well-established RL algorithm. Importantly, the RL learning rate fluctuated according to a signal related to network degree centrality, indexed by dACC activity at the time of witnessing others’ actions, but only insofar as the social observations varied in their informativeness.

The observed dACC response to the degree centrality of the observee relative to that of the observer (i.e., RD) in S2 and S3 is consistent with past evidence showing a key role of this region in facilitating behavioral flexibility and adjusting learning rates for adapting to the external world^41-45^. Our data extend these findings by demonstrating that the regulatory process may also incorporate the topological properties of social connections that underlie learning. In our case, the dACC encoding was seen on an observation-by-observation basis, reflected the degree centrality of the observee and observer simultaneously and independently, existed above and beyond the prediction error and other decision-related variables, and its across-subject variations were predictive of individual differences in the degree modulation effect on behavior.

Importantly, our data also emphasized the absence of a dACC response to the same RD signal in S1 when the network structure was irrelevant to learning. This finding argues against the possibility that the dACC engagement identified in S2 and S3 was due to some low-level visual processing of the network displays, or due to other more general functions of the dACC––such as detecting errors^55^ or monitoring social conflicts^56^––that would be involved across all learning stages nonselectively. Alternatively, the stage-dependent dACC encoding is consistent with a broader proposal of this region, suggested by past neurophysiological and neuroimaging evidence, as representing task-relevant (but not irrelevant) information that supports behavioral changes and guides appropriate action selection^41^. Our data thus suggest the involvement of a high-level, controlled process in evaluating the source of social information in service of learning, and argue against a model of blind, automatic discrimination among social contacts in explaining social information aggregation. More broadly, the observed between-stage differences echo past studies that used data from social media and highlighted the importance of separating information propagation stages, such as those related to the initial transmission and retransmission, in developing mechanistic understandings for rumor dissemination and amplification^26,57^.

While our findings highlighted a role of the dACC specific to S2 and S3, we also observed that, in S1, activity in the PCC, precuneus, and visual cortex correlated with measures of degree centralities. There are several possibilities for how network-related activity in S1 would contribute to learning. One possibility is that the S1 activation is associated with the recognition or representation of network features, which facilitates the flexible usage of those features in the latter stages. Indeed, the loci of the S1 activation were similar to those implicated in representing the internal perception of centralities or other characters of real-life social networks, when subjects were required to view pictures or videos of their acquaintances^29,47,48^. Our data are thus consistent with a possibility that, whereas the degree-modulation effect is stage dependent, some network-related information may be automatically registered in the brain to prepare for the future usage. Alternatively, it is also possible that, S1 activation may reflect, rather than the perception of interpersonal connections, some low-level processing of the network stimuli (e.g., visual processing), which typically shows a more pronounced activation when the stimulus is novel than when the stimulus has been recently processed^58^. To clarify these possibilities and explore how network structure is internally perceived, represented, and transformed into the modulation signal in service of learning, future research is needed to combine the current approach with sociometric methods used for studying real-world connections^59^, to investigate how the brain learns from actual peers without resorting to networks that are artificially structured and displayed.

Degree centrality has long been hypothesized to have a close relationship with social influence in small group interaction and communication^60^. Our finding that the brain modulated learning according to degree-related signals is consistent with two broad accounts previously proposed for how network centrality affects learning. The first has its basis in human and non-human studies that emphasize the role of the structural position in social behavior, suggesting that structurally comparable individuals are facing similar interacting environments, therefore exhibiting similar behavior toward one another^25,28^. In the context of learning, this account proposes that the opportunity to obtain new information through interpersonal interactions may be constrained by one’s location on the interaction network, and can be quantified by location features, such as the degree centrality. A second but not mutually exclusive possibility has its basis in the dynamic nature of network topology: Knowledgeable or successful individuals tend to become highly connected, thus degree centrality may serve to signal an individual’s capability or social status to other individuals^28,61^. Under this possibility, social animals may have evolved to preferentially follow the more “connected” or “prestigious” cospecies, even in controlled experiments where the network structure is fixed and locations are randomized.

Compatible with these diverse lines of proposals, our data additionally highlighted a dual effect of centrality on learning: Higher degree centrality not only amplified one’s social influence, but also reduced one’s susceptibility to others’ influence. This finding is consistent with the behavioral evidence from popular social media, demonstrating that more influential individuals are usually less susceptible to peers’ influence, compared to their less influential counterparts^34^. Our results thus point to an exciting possibility that, while social influence and susceptibility to social influence are often considered as distinct personal attributes^56^, they may be jointly affected by an internal learning system, which approximates the predictive value of others’ information relative to one’s own, in order to cope with the complexity of social environments.

It is worth noting that, owing to the fundamental role of degree centrality in network analyses and its close relationship with a range of network characteristics, we cannot rule out the possibility that alternative network features may contribute to learning. For example, in addition to the degree centrality, which parameterizes the immediate effect of social influence, learning may be affected by measures such as the eigenvector centrality, closeness, clustering coefficient, betweenness, or constraint coefficient, which have been used to examine information propagation from the perspective of long-term, sequential, circular, globally or locally mediating effect, respectively^2^. Our focus on degree centrality reflects the assumption that the brain may be more sensitive to simple, straightforward geometric properties, especially for complex decisions. Indeed, across analyses, there was no evidence that alternative metrics outperformed RD in explaining either the behavioral data or the dACC responses. Future investigation is needed to more firmly isolate and compare the potential influences of various network features at the behavioral and neural levels.

Previous research on the neurocomputational processes of social learning has typically focused on highly simplified interpersonal settings, leaving open whether and how putative RL mechanisms identified in simplistic setups can support behavior in more complex, ecologically-relevant environments. Here, we showed that key features of learning in interconnected contexts were consistent with an error-driven process, similar to those seen in nonnetworked situations. As the network structures examined in the present study were but a sample of immense possibilities of real-world social networks, these results raise questions regarding the scalability and generality of the proposed model. First, the experiment focused on relatively small, 7-node networks, and did not directly speak to larger, more naturalistic settings. Nonetheless, we speculate that, by relying merely on local information, the proposed model may be particularly suitable for scaling up, as individuals in large social groups typically only have access to the local knowledge but not the global information such as the structure of the entire network. Alternative RL algorithms for learning in large-scale networked systems have been developed in control engineering^62^. These algorithms usually aim at optimizing some global network performances (e.g., total reward) and their cognitive and neurobiological feasibility is yet to be evaluated.

Second, the DeGroot-RL model explained choice behavior and task-related neural activity, demonstrating no systematic differences in its explanatory power across networks (Extended Data Fig. 14). Yet, it remains possible that the brain may follow other learning algorithms when facing a different set of networks––for example, deploying Bayesian strategies when making decisions in a line, one of the simplest forms of directed network^37^. Hybrid learning is also possible according to a recent behavioral study suggesting a mixture of Bayesian and DeGroot learning in a relatively more educated (but not less educated) sample^36^. The current study constitutes an initial step toward a neural mechanistic understanding of learning on social networks. Future studies are needed to address whether, and under what circumstances, our findings can be extended to study the potential involvement of multiple learning systems, arbitration between those systems, and individual differences in related processes. Learning on networks offers an excellent opportunity for probing the influence of social structure on the internal tradeoff between computational complexity and learning effectiveness^63,64^.

Social networks have been widely hypothesized to play a key role in many large-scale social phenomena, including vaccine hesitancy, voting behavior, and fake news proliferation, yet the exact mechanisms by which interpersonal connections contribute to these phenomena remain unclear. The current study sheds light on this topic from a neurocognitive perspective, by elucidating how individuals actually experience and interact with a networked environment. Our data provide neural evidence for a bounded rational, network-related filtering of social information, which may result in the spread of misinformation and biased consensus among connected peers. More broadly, this work demonstrates the possibility of developing computationally-tractable and neurobiologically-plausible tools and methods for investigating the complex interplay between social behavior and social embedding in the brain, which may have the potential to translate upward for tackling phenomena in wider society.

## Data availability

Data underlying the findings of this study will be available on the Open Science Framework upon acceptance.

## Code availability

Code supporting the findings of this study will be available on the Open Science Framework upon acceptance.

## Acknowledgements

We thank Yunlu Yin and Yan Wang for assistance with intranet setup and data collection. We also thank the National Center for Protein Sciences and the high-performance computing platform at the Center for Life Sciences at Peking University for facilitating data acquisition and computation. This work is supported by NSFC (32071095 to L.Z.) and Center for Life Sciences at Peking University.

## Author contributions

Y.J. and L.Z. designed the study. Y.J. conducted the experiments. All authors analyzed the data and wrote the manuscript.

## Competing interests

The authors declare no competing interests.

## Notes

### Competing Interest Statement

The authors have declared no competing interest.

